# Dimensionality Reduction with Strong Global Structure Preservation

**DOI:** 10.1101/2025.03.09.642213

**Authors:** Jacob Gildenblat, Jens Pahnke

**Affiliations:** Pahnke Lab, www.pahnkelab.eu; Translational Neurodegeneration Research and Neuropathology Lab, Department of Clinical Medicine, Medical Faculty, University of Oslo, Sognsvannsveien 20, Oslo, NO-0372, Norway; Section of Neuropathology Research, Department of Pathology, Division of Laboratory Medicine, Oslo University Hospital, Sognsvannsveien 20, Oslo, NO-0372, Norway; Institute of Nutritional Medicine, University of Lübeck and University Medical Center Schleswig-Holstein, Ratzeburger Allee 160, Lübeck, D-23538, Germany; Department of Neuromedicine and Neuroscience, The Faculty of Medicine and Life Sciences, University of Latvia, Jelgavas iela 3, Riga, LV-1004, Latvia; Department of Neurobiology, School of Neurobiology, Biochemistry and Biophysics, The Georg S. Wise Faculty of Life Sciences, University of Tel Aviv, G. S. Wise Street, Ramat Aviv, IL-6997801, Israel

**Keywords:** dimensionality reduction, global structure preservation, local structure preservation, MiCS, LMC, visualization

## Abstract

Modern dimensionality reduction (DR) methods, including t-SNE and UMAP, often distort global relationships, limiting the interpretability of embeddings. We introduce two complementary objectives that jointly preserve global geometry and local structure. *Landmark Mantel Correlation* (LMC) aligns high- and low-dimensional distances with respect to a small set of landmarks, providing an efficient global constraint. *Multi-resolution Cluster Supervision* (MiCS) promotes local fidelity by encouraging cluster assignments—estimated across multiple resolutions—to remain predictable after projection. Evaluated on 20 biomedical datasets, UMAP+LMC and MiCS+LMC achieve the best overall performance, demonstrating that global and local structure can be optimized simultaneously rather than being inherently conflicting. Our approach consistently outperforms existing methods for global and local structure preservation, yielding more reliable and interpretable visualizations.

## 1 Introduction

Dimensionality reduction (DR) methods are widely used in data science, both as a pre-processing technique for machine learning and to visualize data by transforming it into 2 or 3 dimensions. These methods can broadly be categorized into those that focus on **global structure preservation** (GSP) and those that focus on **local structure preservation** (LSP). Principal Component Analysis (PCA) [1] and Multi-Dimensional Scaling (MDS) [2] are examples of the former, and t-distributed stochastic neighbor embedding (t-SNE) [3] or Isomap [4] are examples of the latter. Uniform Manifold Approximation and Projection (UMAP) [5] is a widely adopted DR method, with increased scalability and GSP compared to t-SNE.

Despite UMAP’s widespread adoption, particularly in life sciences, several studies have highlighted its limitations, including poor GSP and sensitivity to initialization [6].

Low GSP means that the distances between point clusters and the relationships between points are not guaranteed to be meaningful. Even within clusters, local distances may not reflect true high-dimensional relationships. Modern DR methods, such as UMAP, with high LSP but low GSP, primarily preserve local clusters but do not reliably maintain inter- and intra-cluster relationships—as we demonstrate empirically with extensive benchmarking.

These methods are very useful for data exploration and cluster identification. However, they may be misleading when it is required to distinguish between close points within clusters, analyze global trends in the data, or provide reliable pre-processing for downstream tasks.

The focus of our work is on maximizing GSP while preserving or improving LSP. We offer several contributions that, to the best of our knowledge, were not used for DR before. Extensive benchmarking over 20 biomedical datasets shows that this significantly improves GSP over all methods, including PCA. Our contributions are as follows.

- **Landmark Mantel Correlation (LMC)**: A novel objective that maximizes the correlation of distances from all points to landmarks in both high-dimensional and low-dimensional data, enabling strong global structure preservation.
- **Multi-resolution Cluster Supervision (MiCS)**: A multi-task cluster prediction objective that ensures clusters remain predictable in the reduced space. This simple, interpretable objective, when combined with LMC, provides a competitive balance between LSP and GSP.
- **UMAP+LMC integration**: We demonstrate that LMC can be seamlessly integrated with UMAP, achieving state-of-the-art results in both LSP and GSP.
- **Comprehensive evaluation**: Extensive benchmarking on 20 biomedical datasets reveals that existing methods have poor GSP, while our methods improve performance by a large margin.

## 2 Related work

PCA [7] is a linear dimensionality reduction technique that finds an orthonormal linear projection of high-dimensional data that maximizes the variance. Distances between points along the principal components’ hyperplanes are fully preserved, while for other points, their proximity to the principal components determines their distance preservation. Therefore, PCA is a useful method for high GSP and is often considered a gold standard, although if the selected number of components does not cover the variation in the data, distortions are expected.

UMAP [5] is a widely used non-linear dimensionality reduction technique that constructs a high-dimensional graph of data relationships and embeds it into a lower-dimensional space using a fuzzy topological framework. The focus of UMAP and similar methods is on preserving the local neighborhood of points through the neighbor graph and thus on the LSP. UMAP can preserve some global structure through initialization, for example, by initializing with PCA. Following the formulation recently given in [8], the UMAP objective can be described as

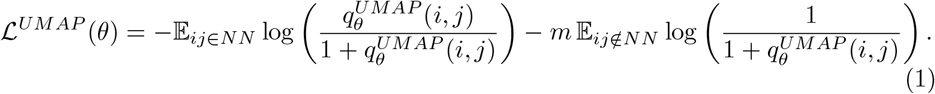

where *q* is the similarity kernel:

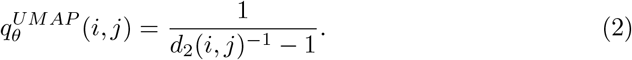

In particular, the fact that *x*_*i*_ is twice as far from *x*_*j*_ as *x*_*k*_ in the original space is lost: UMAP only knows that j is not in the neighborhood set.

PaCMAP [9] is a recent method that follows up on UMAP to explicitly improve GSP. It achieves this through an attraction objective for neighboring pairs of points (like UMAP) and a repulsion objective for distant pairs of points. Repulsing far pairs of points works towards GSP, unlike the emphasis on locality in UMAP. Very recently, LocalMAP [10] extended PaCMAP by dynamically resampling and weighting pairs to focus on local neighborhoods for improved cluster separation.

Our correlation objective uses landmarks to approximate the full correlation while reducing complexity. This approach joins previous DR landmark methods that select a small set of representative points to build embeddings at a fraction of the computational cost [11].

*Landmark Isomap* computes geodesic distances only among the landmarks, then uses them to approximate the global embedding, while *Landmark MDS* runs classical MDS on the landmark set [11, 12]. The remaining points are then placed by simple interpolation from their distances to the landmarks [11, 12].

Regarding our MiCS objective, learning meaningful representations by predicting a fixed set of clusters was proposed in [13], though in the context of representation learning rather than dimensionality reduction.

## 3 Method

### 3.1 The Landmark Mantel Correlation Objective for global structure preservation

The Mantel Distance Correlation metric [14] is used to compare two matrices of distances. It is used for the evaluation of GSP in DR [15] by measuring the correlation of pairs of points in the high-dimensional data and the transformed low-dimensional data. Motivated by this metric, we aim to use it directly as a differentiable objective that we maximize with backpropagation. Since the number of pairs of points is quadratic with the dataset size, we approximate the objective with a set of L landmarks. In all our experiments, we sample the landmarks uniformly.

Let *D*_*X*_, *D*_*Y*_ *∈* ℝ ^*N ×L*^ be the matrices of distances from *N* points to a set of *L* landmarks *ℒ*, with entries

Let vec be the column stacking operator that flattens a matrix into a vector. The LMC is the cosine similarity between the mean-centered and normalized distance profiles:

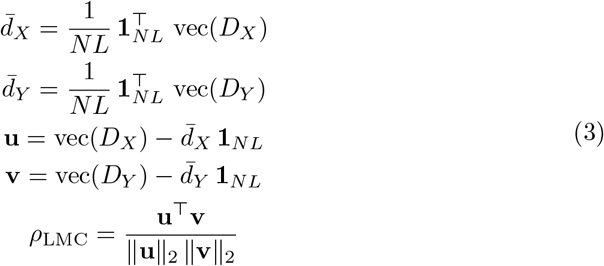

It is helpful to examine the gradients of the LMC objective to understand how points are positioned. The gradient with respect to the low-dimensional distances *v* is:

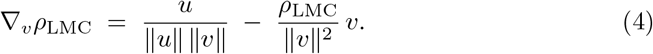

This gradient reveals the dynamics of attraction and repulsion:

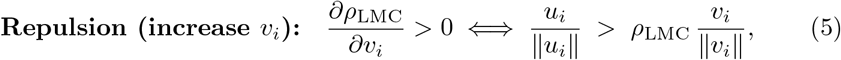

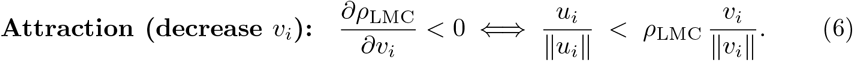

Unlike methods such as UMAP and PaCMAP — where a pair’s update is essentially fixed (attractive for graph edges, repulsive for sampled non-edges) — our LMC objective induces *globally coupled* forces. The attraction/repulsion for any pair depends on the entire configuration via the global distance profiles (point–to–landmark distances): as the embedding evolves, a given pair can switch from attraction to repulsion (and vice versa), reflecting changes in the global arrangement rather than a preassigned edge/non-edge role.

Although LMC is driven by global alignment, its forces also act on nearby pairs and thus can *enhance local structure*. In practice, we find that adding LMC to a baseline (e.g., UMAP or MiCS) consistently improves local-structure metrics while substantially strengthening global-structure preservation.

A visual example of LMC on synthetic datasets is given in Figure 1. LMC and PCA are able to preserve the global structure, unlike modern DR methods. In our quantitative evaluations, we show that LMC significantly outperforms PCA.

**Fig. 1.**
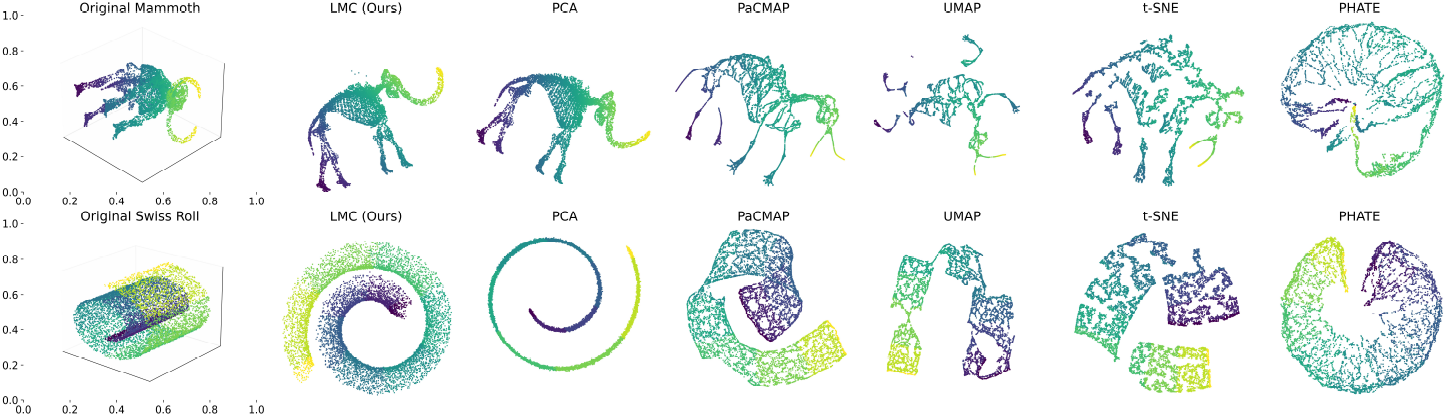
Comparing dimensionality reduction methods for global structure preservation on synthetic toy datasets. Our proposed Mantel Landmark Correlation objective and PCA can preserve the global structure, while modern DR methods like UMAP sacrifice the global structure for the local structure.

**Fig. 2.**
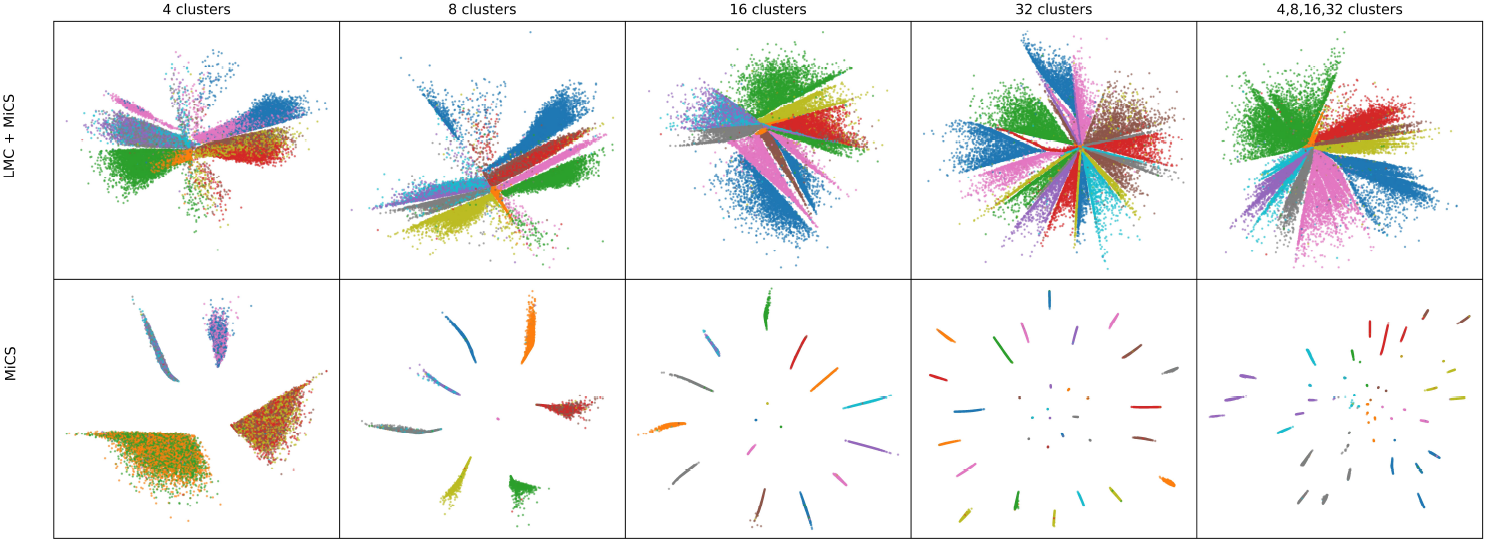
MNIST [16] embeddings colored by digit labels for the proposed Landmark Mantel Correlation (LMC) + Multi-Resolution Cluster Supervision (MiCS) under different clustering combinations. Using a multi-task loss over several clustering sets (right) improves local grouping by pulling together points with the same label that single-*K* tasks may split. The red clusters, for example, are reoriented near one another.

### 3.2 Multi-Resolution Cluster Supervision (MiCS) for local structure preservation

DR methods with strong LSP are often used to visualize or detect clusters; in 2D scatter plots, points are typically colorized by cluster assignments. We optimize for this *explicitly* to enforce cluster visibility in the low-dimensional space.

A smaller number of clusters corresponds to a *coarse* resolution (each cluster covers a larger portion of the data and represents broad neighborhoods). A larger number of clusters corresponds to a *fine* resolution (smaller neighborhoods). We hypothesize that preserving multiple resolutions in the visualization is beneficial: the embedding should reveal fine-scale clusters while coherently arranging them into higher-level groups.

To make clusters at different resolutions visible and to reduce clustering noise, we optimize a *multi-task* objective over multiple sets of cluster labels. Concretely, for each clustering, we learn a linear classifier (with bias) on the low-dimensional embeddings, and use a temperature parameter *T >* 0 for stable, faster convergence. The classifiers and the embedding are co-learned: clusters must be separable in the low-dimensional space for the linear heads to classify points successfully. The MiCS objective is given in Eq. (7).

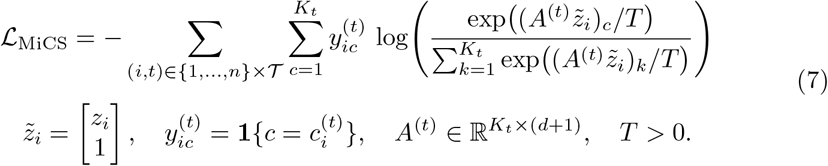

Clustering can be obtained by any clustering algorithm, or alternatively by supervised labels.

### 3.3 Combining the local and global objectives

Deviating from UMAP/t-SNE, we initialize the embedding with i.i.d. Gaussian noise (not PCA). The LMC correlation *ρ*_LMC_ *∈* [*−*1, 1] may be negative at early iterations. To emphasize learning when alignment is poor while keeping optimization stable, we dynamically weight the correlation term by the inverse of the correlation.

The loss is then

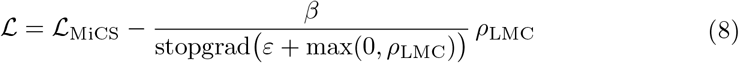

Here, stopgrad(*·*) denotes a detached quantity (implemented as.`detach()` in PyTorch), and the max(0, *ρ*_LMC_) clamps negative correlations before inversion.

For MiCS, we use a multi-task objective, predicting all clusters, with k = 4, 8, 16, 32.

### 3.4 Computational complexity and balancing the loss terms

The LMC objective requires storing the distances of all points from L landmarks, and computing distances from all N data points to the L landmarks. Therefore, the complexity is O(NL) in both storage and computation. The MiCS objective, on the other hand, requires simply classifying each of the N points with the linear classifiers and is much faster.

To condition the embedding on local structure preservation earlier in the optimization, as well as to speed up the computation, the LMC loss weight is dynamically varied. The LMC loss weight decays every T epochs, starting from T = 50, with a decay factor of 0.95, until it is finally applied every epoch towards the end of the optimization.

### 3.5 Combining UMAP and the Landmark Mantel Correlation

Similarly, we plug the LMC objective into the UMAP loss given in equation (1) and optimize for both

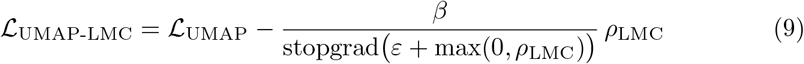

We found it beneficial to start the LMC objective after several epochs. Empirically, if LMC is applied from start, we get lower convergence. See the hyperparameter examples in Figure 8.

## 4 Experiments

We conducted a comprehensive evaluation of our approach against eight state-of-the-art DR methods. The benchmarking was carried out on twenty diverse biomedical datasets, using ten random seeds to ensure robustness to initialization and reproducibility. Figure 3 illustrates the different methods for scRNA datasets from the evaluation. Performance was assessed on complementary quantitative metrics, providing a balanced view of both local and global structure preservation. The next sections describe the DR methods we compared against, the metrics, and the datasets.

**Fig. 3.**
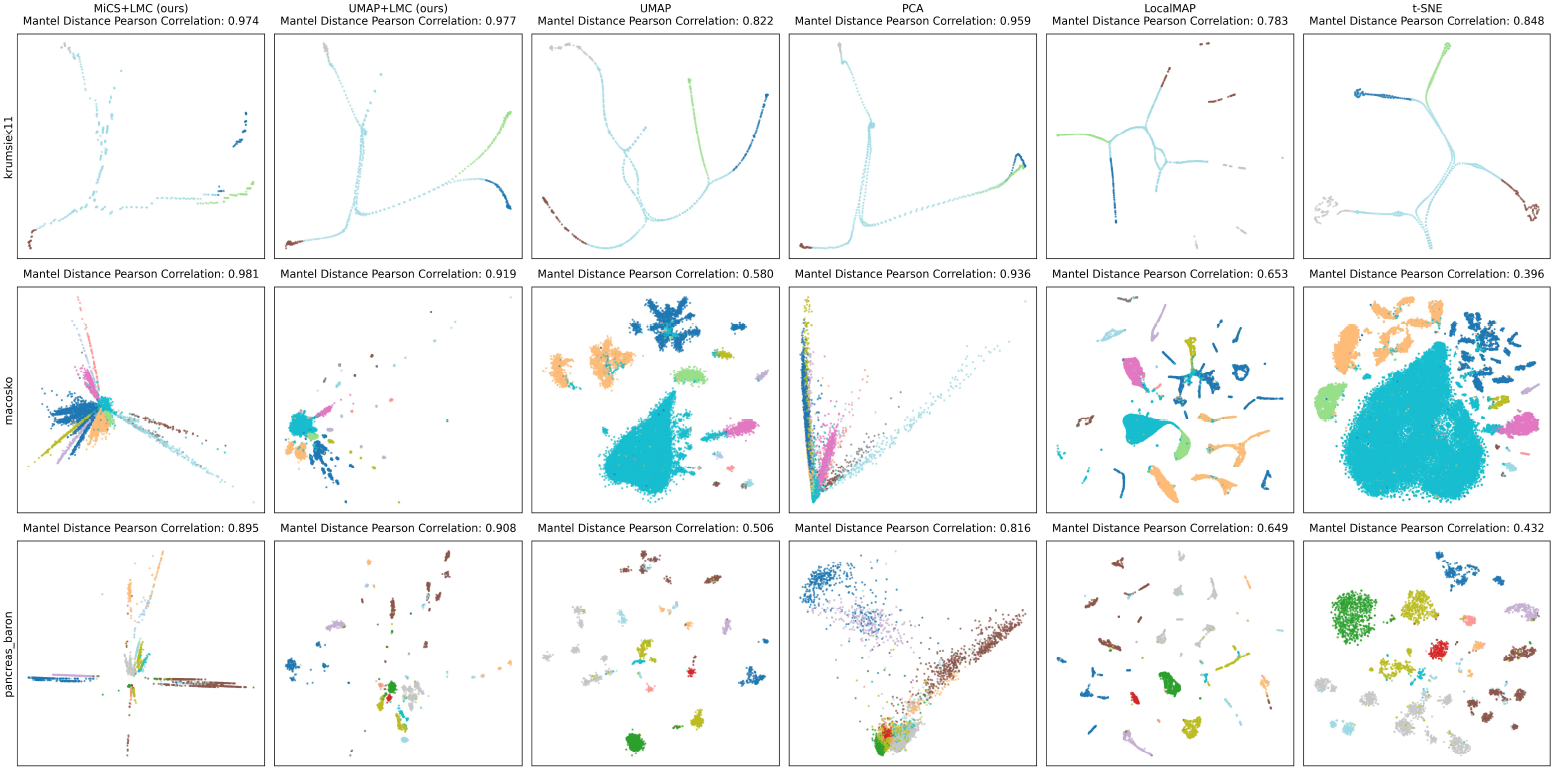
Examples on scRNA datasets from the evaluation benchmark. Our proposed methods achieve high global structure preservation while maintaining local information that is lost in PCA. The LMC can be combined with any DR method to benefit from improved global and local structure.

### 4.1 Evaluated Dimensionality Reduction Methods

We evaluated our two proposed methods (MiCS+LMC, UMAP+LMC) against eight state-of-the-art DR methods, including two recently introduced within the past year. We chose PCA [1] as a widely used method that excels in GSP, and the rest are modern, recent methods, with t-SNE [3] and UMAP [5] being very widely used DR methods for LSP. A summary of the DR methods is given in Table 1.

**Table 1.** Benchmarked dimensionality-reduction methods, with brief strategies and emphasis on local vs. global structure.

| Method | Year | One-line description | Focus |
| --- | --- | --- | --- |
| LocalMAP [10] | 2025 | Locally adjusted graph that dynamically refines neighborhoods to separate true clusters. | Global/Local |
| DTNE [17] | 2025 | Diffusive topology-preserving manifold distances for single-cell embeddings. | Global/Local |
| PaCMAP [9] | 2021 | Schedules near, mid-near, and far pairs to balance local and global geometry. | Global/Local |
| PHATE [18] | 2019 | Diffusion operator and “potential distances” for multi-scale manifold structure. | Global/Local |
| TriMap [19] | 2019 | Triplet constraints with near positives and many far negatives to maintain large-scale relations. | Global/Local |
| UMAP [5] | 2018 | Fuzzy kNN graph + cross-entropy; attractive edges with negative-sampling repulsion. | Local |
| t-SNE [3] | 2008 | KL divergence between high/low affinities with strong non-neighbor repulsion. | Local |
| PCA [1] | 1901 | Linear projection maximizing variance (orthogonal components). Very strong global structure preservation. | Global |

### 4.2 Metrics

We use a total of seven metrics, four for local structure preservation and three for global structure preservation.

- **Trustworthiness** (local) [20]. Penalizes *intrusions*—points that become *k*-NN neighbors in the embedding *Y* but were not *k*-NN neighbors in the original space *X*.
- **Continuity** (local). Penalizes *exclusions* —true *k*-NN neighbors in *X* that are no longer *k*-NN neighbors in the lower-dimensional data *Y*.
- **MRRE Intrusions** (local) [21]. Measures *intrusions* using relative rank errors within a neighborhood of size *k*.
- **MRRE Exclusions** (local) [21]. Measures *exclusions* using relative rank errors within a neighborhood of size *k*.
- **Pearson Mantel correlation** (global) [14, 15]. Correlation between the vectorized pairwise *distance* matrices in *X* and *Y*.
- **Spearman Mantel correlation** (global) [14, 15]. Rank-based correlation between the vectorized pairwise *distance* matrices in *X* and *Y*.
- **Centroid-level Mantel correlation** (global). Following [8], cluster the low-dimensional embedding at *K* ∈ {4, 8, 16, 32}; compute inter-cluster *centroid* distance matrices in *X* and *Y*, and report their Mantel Pearson correlation.

### 4.3 Evaluation datasets

We evaluated 20 datasets spanning diverse biomedical modalities, detailed in Table 2. These included ten single-cell RNA sequencing datasets (e.g., retina, PBMCs, pancreas, hematopoiesis, and neurogenesis), three spatial transcriptomics datasets from 10x Visium, and one mass spectrometry imaging dataset of the mouse brain [22]. Beyond transcriptomics, we incorporated proteomics (Mice protein), a high-dimensional cancer expression challenge dataset (Arcene), and a synthetic generegulatory network (Krumsiek11). Finally, four classical biomedical benchmarks were included: Breast Cancer WDBC, QSAR biodegradation, Parkinson’s voice, and Low birth weight.

**Table 2.** Biomedical datasets used in our evaluation. scRNA-seq = single-cell RNA sequencing; ST = spatial transcriptomics; MSI = mass spectrometry imaging.

| Dataset | Modality | Description |
| --- | --- | --- |
| Macosko 2015 [23] | scRNA-seq | Mouse retina Drop-seq cells; foundational large-scale single-cell dataset. |
| Zheng PBMC 68k [24] | scRNA-seq | Human peripheral blood mononuclear cells; canonical benchmark reference. |
| Paul15 [25] | scRNA-seq | Mouse myeloid progenitors across hematopoietic differentiation. |
| PBMC3k (processed) [26] | scRNA-seq | 10x PBMC tutorial dataset widely used in Scanpy examples. |
| PBMC68k (reduced) [24, 27] | scRNA-seq | Subsampled 68k PBMCs for faster benchmarking runs. |
| Pancreas (Muraro) [28] | scRNA-seq | Human pancreas single cells spanning major endocrine and exocrine types. |
| Pancreas (Baron) [29] | scRNA-seq | Human/mouse pancreas cells from multiple donors with rich annotations. |
| Pancreas (Segerstolpe) [30] | scRNA-seq | Human islet cells, including healthy and type 2 diabetes samples. |
| Visium Human Lymph Node [31] | ST (Visium) | Spatial transcriptomics of human lymph node on 10x Visium. |
| Krumsiek11 GRN toy [32] | synthetic GRN | Simulated gene-regulatory network modeling myeloid differentiation. |
| Pancreas (scVelo) [33] | scRNA-seq | Pancreatic endocrinogenesis trajectory used in RNA velocity demos. |
| Dentate gyrus neurogenesis [34] | scRNA-seq | Mouse hippocampal cells capturing postnatal neurogenesis trajectory. |
| Squidpy Visium H&E demo [35, 36] | ST (Visium, H&E) | Example H&E-stained Visium dataset bundled with Squidpy. |
| Mice Protein (OpenML) [37] | proteomics | Protein expression profiles from a mouse trisomy study. |
| Arcene (OpenML) [38] | expression | Cancer vs. control samples with high-dimensional features (FS challenge). |
| Breast Cancer WDBC [39] | clinical | Diagnostic features from digitized breast mass FNA images. |
| QSAR biodegradation [40] | chemical | Molecular descriptors labeled as ready/not-ready biodegradable. |
| Parkinson’s voice [41, 42] | audio (clinical) | Acoustic features for the detection of Parkinson’s disease. |
| Low birthweight (birthwt) [43] | clinical | Maternal/infant variables associated with infant birthweight. |
| Mouse brain MSI [22, 44] | MSI | Mouse brain lipidomics mass spectrometry imaging scans with 5,005 features. |

## 5 Results

### 5.1 Average performance across 20 datasets: UMAP+LMC and MiCS+LMC lead on global and local structure

Figure 4 summarizes results by first averaging over seeds within each dataset, then aggregating across the 20 datasets (error bars: std across datasets). On average, both **UMAP+LMC** and **MiCS+LMC** lead on most metrics. Crucially, they deliver strong *global* fidelity, exceeding PCA and prior baselines, while maintaining *local* neighborhood quality (LSP). In particular for GSP (Mantel–Pearson), UMAP+LMC increases the average from *≈* 0.49 (UMAP) to ≳ 0.92.

**Fig. 4.**
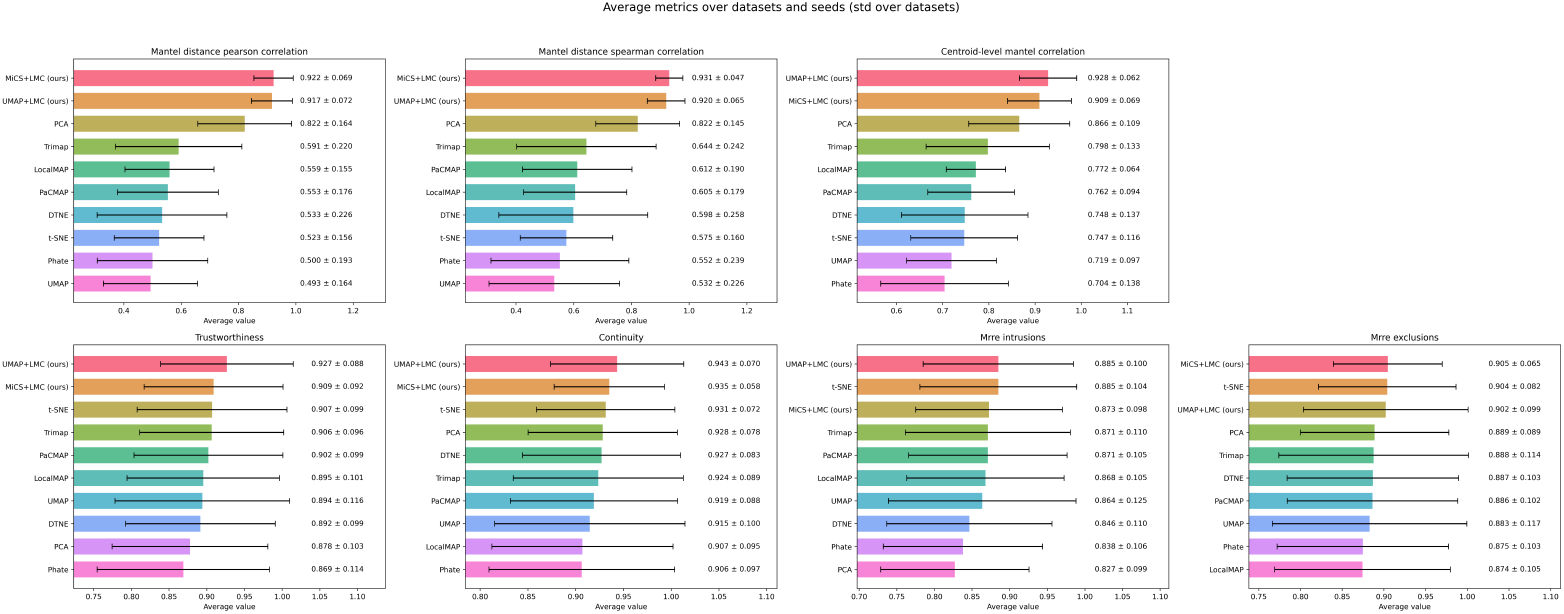
Average performance across datasets. For each dataset, metrics are averaged over 10 random seeds; points show the mean across 20 biomedical datasets, and error bars indicate the standard deviation across datasets. Our methods obtain the best average results on most metrics and substantially close the global-structure preservation (GSP) gap (Mantel–Pearson improves from UMAP’s 0.49 to 0.92), while remaining competitive on local-structure preservation (LSP).

### 5.2 UMAP+LMC: best per-dataset trade-off between global and local structure preservation

Figure 5 reports per-dataset scores (mean *±* std across seeds) and aggregates them via average rank and win counts. **UMAP+LMC** and **MiCS+LMC** both achieve strong GSP while retaining competitive LSP. Compared to prior baselines (excluding MiCS+LMC), **UMAP+LMC** yields the best overall average rank and the highest number of strict wins, indicating the most balanced performance when both local and global fidelity matter. When global structure is prioritized, **MiCS+LMC** provides a favorable trade-off and remains competitive on LSP.

**Fig. 5.**
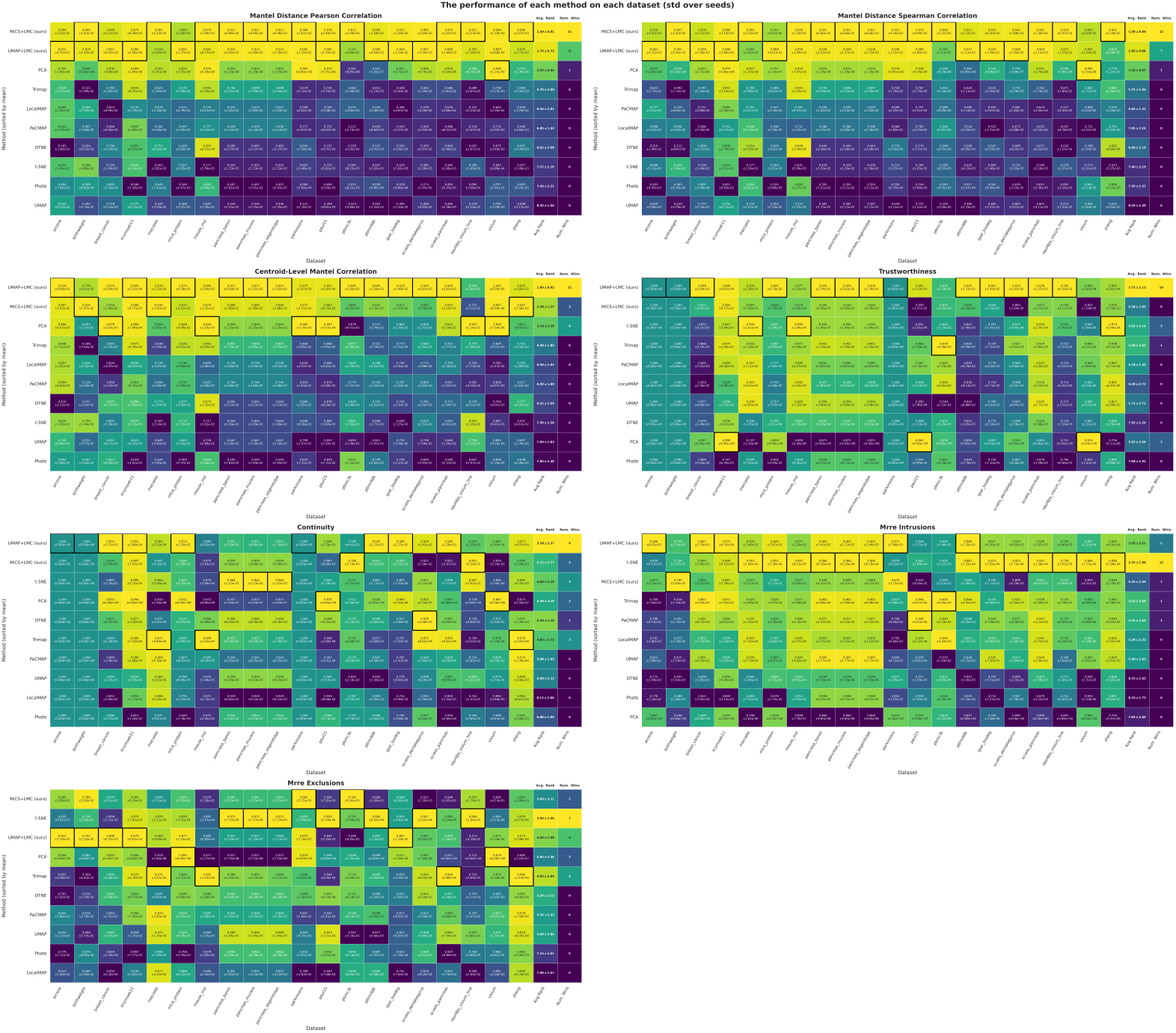
Performance over individual datasets. Averaged over ten random seeds, with standard deviations computed over the ten random seeds. The highest performing method in each dataset is highlighted with a black border. The two last columns show the average rank of each method across the datasets (lower is better, std across datasets) and the number of times each method was ranked first. Our proposed methods significantly improve the global structure. Overall our UMAP+LMC performs best in terms of ranks and number of wins, and is the most robust method.

### 5.3 Ablations and hyperparameter sensitivity

We also studied the effect of hyperparameters for UMAP+LMC and MiCS+LMC. Figure 6 summarizes results across 20 datasets when enabling/disabling the LMC term and the auxiliary objectives. LMC is the primary driver of high *global-structure preservation (GSP)*; however, it benefits from being combined with a complementary objective. When paired with either MiCS or UMAP, LMC preserves or improves *local-structure preservation (LSP)* while dramatically increasing GSP. By contrast, MiCS alone does not yield high GSP, but the combination MiCS+LMC improves both LSP and GSP.

**Fig. 6.**
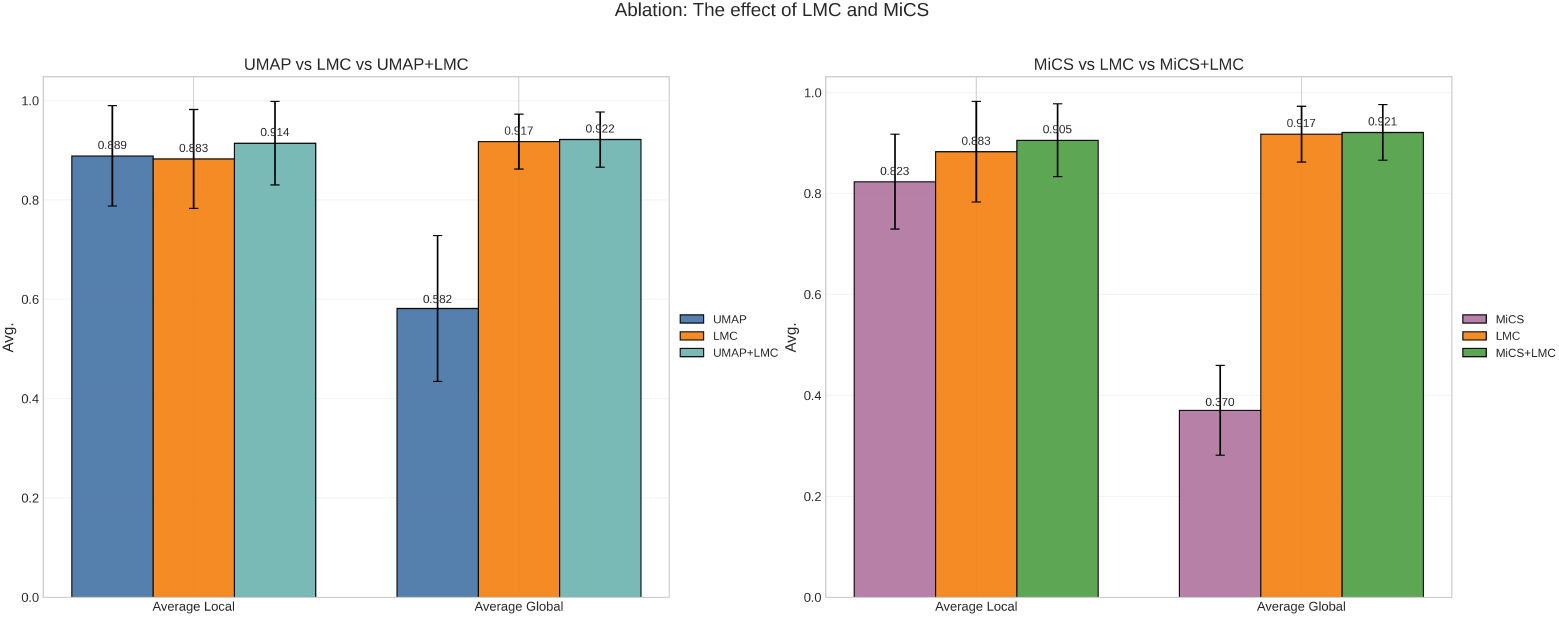
Ablation of MiCS and LMC objectives over the 20 evaluation datasets. LMC alone attains high global-structure preservation (GSP) but lower local-structure preservation (LSP) than when combined with either MiCS or UMAP. MiCS alone does not yield high LSP, but is synergistic with LMC. Overall, pairing LMC with UMAP or MiCS achieves high GSP while maintaining strong LSP.

Figures 7 and 8 provide visual sensitivity plots on `fashion-mnist` [45] for MiCS+LMC and UMAP+LMC, respectively. For most hyperparameters there is a broad operating range where performance is near-optimal; increasing them beyond that range typically leads to saturation and, in some cases, mild degradation. In Figure 7, for example, expanding the number of clusters used by the multi-task MiCS objective steadily improves performance but saturates around *k* = 128.

**Fig. 7.**
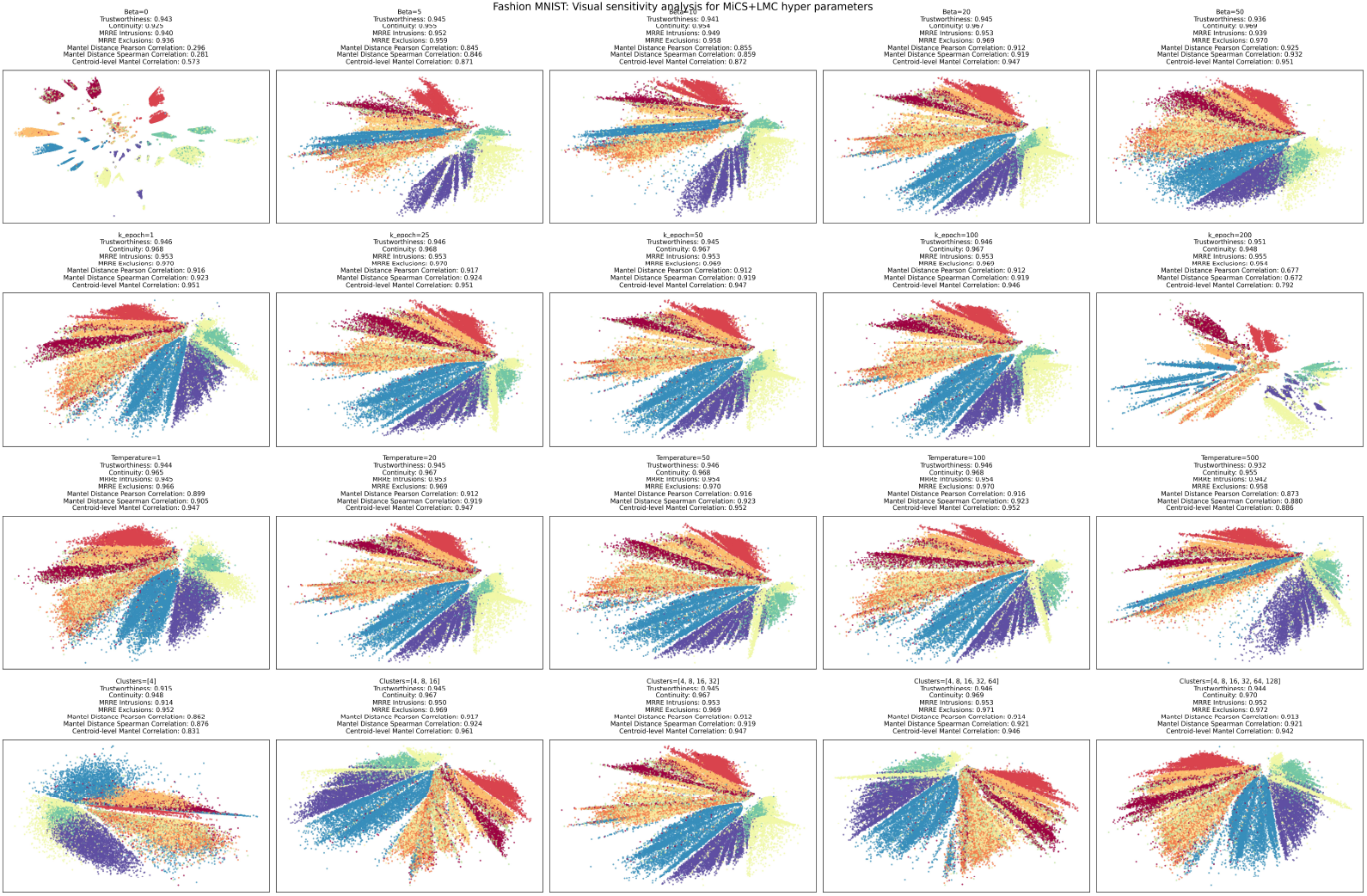
Visual comparison of different hyperparameters for MiCS+LMC on the Fashion-MNIST dataset (not used in the evaluation).

**Fig. 8.**
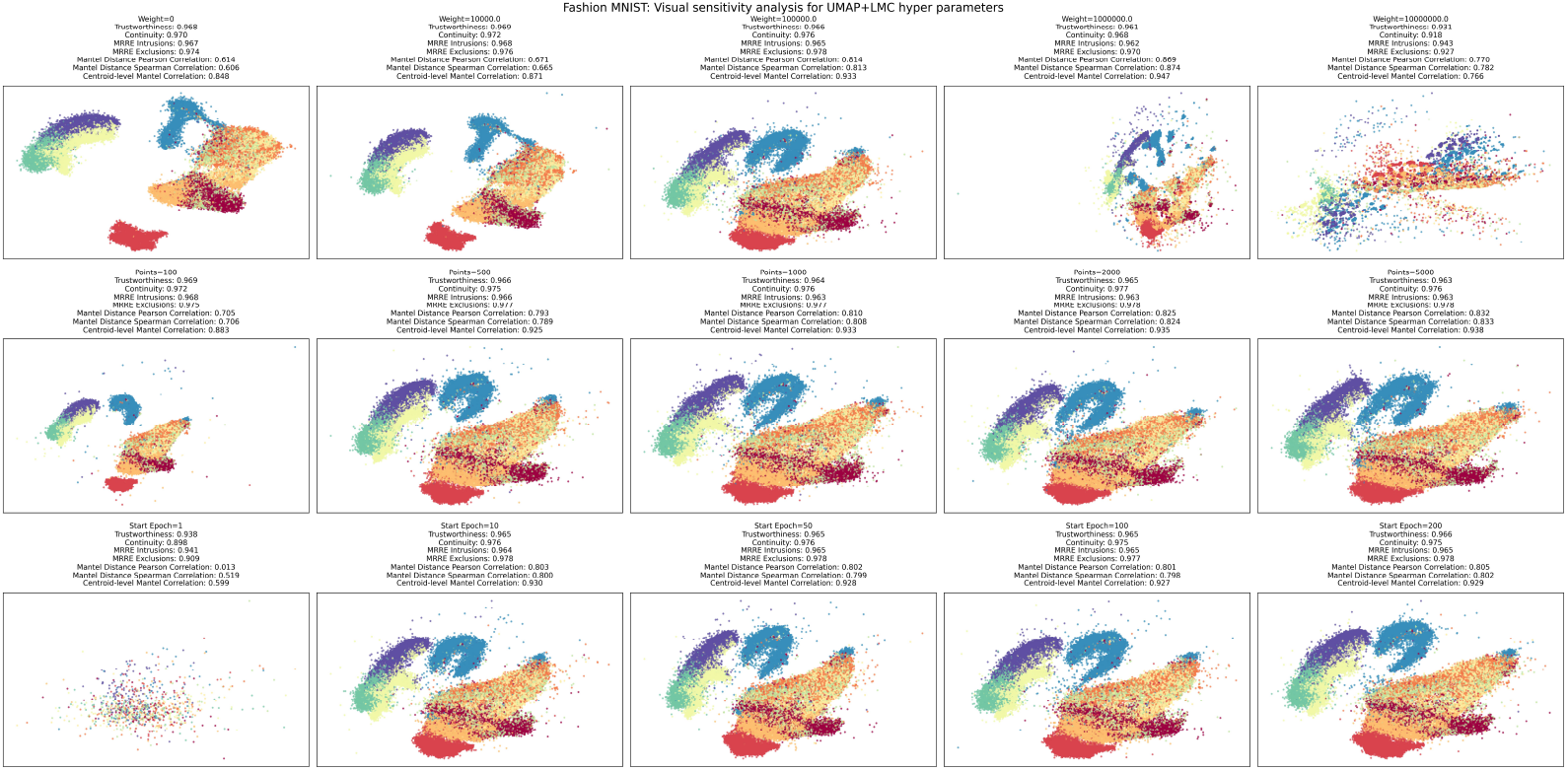
Visual comparison of different hyperparameters for UMAP+LMC on the Fashion-MNIST dataset (not used in the evaluation). The top left image with Weight=0 is UMAP alone without any addition.

In both families of experiments, the number of landmarks exhibits clear diminishing returns: beyond *K ≈* 500 landmarks, both LSP and GSP metrics change very little.

### 5.4 Summary of details for reproducibility, and hyperparameter fairness

Unless otherwise specified, MiCS+LMC was configured with 1000 landmark points, *β* = 20, *T* = 20, and *k*_*epochs*_ = 50. For UMAP+LMC, we set *β* = 90, 000 and applied LMC after 10 epochs. These hyperparameters were tuned during development using datasets not included in the evaluation, primarily MNIST, Fashion-MNIST, and in-house Mass Spectrometry Imaging datasets.

For MiCS+LMC, in all experiments, we used UMAP to reduce the high-dimensional data into an intermediate-dimensional representation with 10 dimensions, and then clustered the intermediate representation.

For all other methods, we use the default hyperparameters. For TriMAP, we modified the source code to accept a random seed.

## 6 Discussion

This work addresses the fundamental trade-off between local and global structure preservation in dimensionality reduction. Our Landmark Mantel Correlation (LMC) objective demonstrates that global structure preservation can be substantially improved through landmark-based correlation maximization, while our Multi-resolution Cluster Supervision (MiCS) approach provides a simple method for local structure preservation.

The key insight is that landmark-based approaches can capture global relationships that are inherently lost in graph-based methods like UMAP and t-SNE. Traditional graph-based methods construct local neighborhoods and optimize pairwise relationships within these neighborhoods, which naturally emphasizes local structure at the expense of global geometry. In contrast, landmark-based correlation optimization considers the entire data manifold simultaneously by comparing distance profiles to reference points, enabling the preservation of both fine-grained local relationships and coarse-grained global structure. By maximizing correlations between high- and low-dimensional distance profiles to reference points, LMC achieves GSP performance exceeding even PCA, traditionally considered the gold standard for global structure preservation. LMC’s gradient reveals that it creates dynamic forces that depend on the global arrangement of points, unlike the static neighbor-based forces in existing methods.

This approach has particular significance for biomedical data analysis, where both local clustering (e.g., cell type identification) and global relationships (e.g., developmental trajectories, disease progression) are crucial for interpretation. Our methods enable researchers to visualize complex biological processes while maintaining confidence in both the local neighborhood structure and the global organization of their data.

Our evaluation in biomedical datasets shows dramatic improvements in GSP metrics (Mantel-Pearson correlation increased from 0.49 to 0.92) while maintaining competitive LSP performance. This represents a substantial advance over recent methods like LocalMAP [10] and DTNE [17], which focus on improving local neighborhoods through dynamic resampling or diffusive distances but still struggle with global structure preservation. Our landmark-based approach fundamentally differs by optimizing global distance correlations rather than refining local graph structures, enabling simultaneous optimization of both local and global structure preservation.

Ablation studies demonstrate that LMC and MiCS are complementary: LMC alone achieves excellent GSP but limited LSP, while MiCS alone provides moderate LSP improvements. Their combination yields the best overall performance, indicating that global and local structure preservation can be optimized simultaneously rather than being inherently conflicting objectives.

The computational efficiency of our approach—O(NL) for LMC and the linear complexity for MiCS—makes it scalable to large datasets. The landmark sampling strategy provides a natural way to balance computational cost with representation quality, and our results suggest that relatively small numbers of landmarks (1000 in our experiments) can capture essential global structure. This efficiency enables practical application to modern large-scale datasets. To ensure reproducibility and facilitate adoption, our implementation is freely available as open-source software with comprehensive documentation and example workflows.

Practically, our methods are particularly valuable for exploratory data analysis scenarios where researchers need to understand both local clustering patterns and global data organization. For example, in single-cell RNA sequencing analysis, researchers often need to identify cell types (local structure) while understanding developmental trajectories or disease progression (global structure). Similarly, in mass spectrometry imaging, both local metabolite co-localization and global tissue organization patterns are important for biological interpretation.

### 6.1 Limitations

Several limitations should be acknowledged. The landmark selection strategy (uniform random sampling) may not be optimal for all datasets, particularly those with highly non-uniform density distributions. The MiCS objective relies on the quality of the reference clustering, which could be improved through more sophisticated clustering strategies. Additionally, our evaluation focused on biomedical datasets; performance on other domains remains to be validated. Another limitation is that the landmarks used are constant. This means that a poor choice of landmarks would effect all optimization iterations. In addition, as the ablation has shown, the MiCS objective by itself does not provide high LSP, and is rather used as a complementary objective to LMC.

### 6.2 Future Work

Future work could explore adaptive landmark selection strategies that consider data density or manifold curvature, dynamically weighting pairs in the correlation objective based on their importance for global structure, and extension of MiCS to incorporate additional supervision signals such as temporal information or known biological pathways. If data labels are included in the MiCS objective, it provides a trivial form to perform supervised dimensionality reduction. MiCS can be particularly adapted for biomedical imaging visualization, where clustering is commonly used for visualization by coloring pixels based on their assigned cluster. MiCS allows superimposing several clustering results on a single summary imaging.

An exciting direction involves developing parametric versions of our methods that can learn to embed new, unseen data points [46]. By training neural networks to approximate the MiCS+LMC embedding function, we could enable out-of-sample extension. Furthermore, the MiCS+LMC framework could serve as a powerful self-supervision signal for representation learning, where the learned embeddings provide high-quality targets for training more complex models on downstream tasks.

Beyond cluster-based supervision, future work could explore alternative supervision signals such as soft distances from cluster centroids, or manifold-based geometric properties. These approaches could provide more nuanced supervision than hard cluster assignments while maintaining the computational efficiency and interpretability of our current framework.

The broader implications of this work extend beyond dimensionality reduction to the fundamental challenge of maintaining interpretability in high-dimensional data analysis. As datasets continue to grow in size and complexity, methods that can preserve both local and global structure will become increasingly important for ensuring that computational analyses remain biologically meaningful and scientifically interpretable.

### 6.3 Conclusion

This work demonstrates that the local-global structure preservation trade-off in dimensionality reduction can be substantially improved through landmark-based correlation optimization. Our methods provide valuable tools for pattern analysis applications where interpretable, faithful data representations are essential, particularly in biomedical research where both local clustering and global organization are crucial for scientific insight.

## Supporting information

R1 - answers to reviewer comments

## Code availability

The code is available on GitHub (github.com/jacobgil/pcc).

## Author contributions

J.G. and J.P. planned and performed the investigation, wrote and revised the manuscript.

