## Supplementary material for "Dimensionality Reduction with Strong Global Structure Preservation": R1 - answers to reviewer comments

**Letter to the editor**

Following the feedback, **we have performed a major revision and completely re-written** the manuscript, simplified and focused it.

- We extensively strengthened  the quantitative evaluation. We extended the evaluation to be performed on 20 bio medical datasets. This is now one of the largest quantitative evaluations of DR methods for biomedical applications.
- The evaluations are now done over 10 seeds, and we report error bars in the quantitative evaluation.
- We re-wrote the introduction.
- The related work section was extended to include more work, as well as more detail about related work, relevant to our comparison to them.
- We add motivations for our methods, elaborate on the rational and connect to previous work.
- The methods themselves have been improved since the submission.
- For improved clarity and focus, the methods were renamed.
- We added a section for ablations, and hyper parameter sensitivity.
- We added a dedicated section discussing details for reproducibility.
- We improved the visual examples, and added more visual examples that illustrate the methods, in addition to the extended quantitative evaluation.
- We also adapted the title of the work to fit better to the current content after revision.

**Reviewer 1**

This manuscript proposes a new dimensionality reduction method, PCC, and claims that it significantly outperforms existing approaches in preserving global structure. However, the paper suffers from several critical issues that undermine its academic contribution:

1. **Lack of formal derivation and theoretical analysis**: The methods section provides only surface-level definitions of the loss functions without systematic mathematical derivation or theoretical justification. Key aspects such as the differentiability of the Spearman rank approximation, the impact of reference point sampling on computational complexity and convergence, and the rationale behind the local structure preservation objective are not rigorously derived, leaving the methodological foundation weak.

   We thank the reviewer for this valuable feedback. We removed the Spearman rank correlation objective. We added reasoning for the clustering objective, and now discuss attraction/repulsion properties of the correlation objective. We now also discuss computational complexity.

2. **Insufficient quantitative evaluation**: While some visual comparisons are presented, most results remain qualitative. The manuscript lacks comprehensive quantitative tables or statistical tests, and even contains placeholders such as “Table ??,” which suggests incomplete or careless preparation. Without robust numerical comparisons, the conclusions are unconvincing.

   Following this feedback, we added **extensive** quantitative evaluation. We extended the evaluation to focus on **20 biomedical datasets**. We now report **7 quantitative metrics for 10 seeds across 20 bio medical datasets**. We added several figures that show average and ranking performance of our methods over the metrics. We report error bars over the 10 random seeds, and also error bars over the 20 datasets.

3. **Overall quality issues**: The manuscript contains numerous flaws in formatting, references, and experimental details (e.g., the misspelling of “Introdution,” incomplete reference entries), which further weaken the integrity of the submission.

   Thank you for this valuable feedback. We have fully rewritten the manuscript, fixed references, grammar, as well as tried to improve the clarity.

4. **Unclear distinction from existing methods**: Although the authors claim that PCC substantially outperforms methods such as UMAP and PaCMAP, there is no clear theoretical or mathematical comparison. Moreover, the experiments do not discuss hyperparameter fairness, making it unclear whether the reported improvements are due to the method itself or differences in experimental design.

   Following this valuable feedback, we now discuss, both in the related work section and in the method section, the differences between UMAP and PaCMAP, including motivating the correlation objective as a dynamic attraction and repulsion loss.

For hyper parameters, we added clarifications as well as a sensitivity analysis section.

5. **Lack of quantitative validation in applications**: In the biomedical imaging experiments, only qualitative visualizations are provided. No objective metrics (e.g., clustering scores, classification accuracy, alignment with pathological annotations) are reported, leaving the practical value of PCC unsubstantiated.

   Following this feedback we removed the imaging experiment that was intended as a visual example. Instead **we increased the number of benchmarked datasets from 9 to 20, focusing on biomedical datasets**, where practitioners extensively use dimensionality reduction for visualization.

6. **Absence of ablation studies**: PCC is composed of both a global correlation objective and a local cluster separability objective. However, the manuscript does not include ablation experiments to validate the independent contribution of each component, making the interpretation of the method's effectiveness incomplete.

   We thank the reviewer for this feedback. Following this, **we added a detailed section on ablation and hyper parameter sensitivity**.

7. **Code and reproducibility concerns**: Although the authors state that code is available on GitHub, the main text omits crucial implementation details (e.g., how the number of reference points is chosen, the principle behind setting parameter $\beta$). As a result, readers cannot reliably reproduce the results based solely on the paper.

   Following this feedback, **we added a section on details needed for reproducibility**, and clarifications on how the parameters were chosen. The new hyper parameters sensitivity section, added following this valuable feedback, demonstrates the rationale for the hyper-parameter selection both visually and quantitatively.

**Reviewer 2**

This manuscript introduces a novel dimensionality reduction method, termed Clustering with Correlation Conservation (PCC), which aims to achieve state-of-the-art performance in global structure (GS) retention while maintaining competitive local structure (LS) preservation. However, several aspects of the manuscript require further clarification and improvement before it can be considered for publication.

1. The complete name of any method mentioned in the paper should be provided upon its first occurrence. Furthermore, while the authors state in the abstract that 'We quantitatively benchmark PCC against existing methods and demonstrate its utility in medical imaging', no relevant discussion pertaining to 'medical imaging' is presented in either the introduction or the previous work section.

   Thank you for this valuable feedback. To simplify, we removed the medical imaging visual example, and extended the quantitative evaluation from 9 datasets to focus instead on 20 biomedical datasets. We also now introduced the complete method names as suggested.

2. It is unclear why the directly employs average weighting. The authors are encouraged to provide a detailed explanation, including theoretical or empirical justification. At present, this term appears to be primarily a straightforward combination of existing methods, the authors should clearly articulate the key novelty.

   We removed the Spearman rank correlation objective in order to simplify and focus the manuscript. Following this valuable feedback we tried clarifying the novelty and the rational of each objective.

3. The comparison methods in the experiment section should incorporate the most recent DR methods to enhance the novelty of the proposed method.

   We thank the reviewer for this feedback. Following this, we added two strong dimensionality reduction methods from 2025, LocalMAP (a continuation of PaCMAP) and DTNE.

4. The sensitivity of the parameters used in the PCC method should be discussed.

   Following this feedback we added a **dedicated section on Ablations** and hyperparameter sensitivity, both visual and quantitative.

5. The concept of clustering is referenced in both the title and keywords. In the experiment section, the author should incorporate an analysis of the PCC method using clustering metrics.

   Thank you for this valuable feedback. We performed a major update of the paper, including changing method names and the title to improve the focus and clarity. Following this feedback we added one global structure preservation metric that utilizes clustering - the mantel distance correlation between cluster centers, averaged over several choices of number of clusters.

6. The article contains certain formatting issues, such as ‘Table ??’ on page 9 and ‘???’ within [2] on page 12. The author should carefully review the formatting of the article once more.

   We thank the reviewer for this feedback and hope we substantially improved the article formatting.

7. The author should refer to relevant DR methods within the last two years.

   We thank the reviewer for this feedback. Following this we added two strong dimensionality reduction methods from 2025, LocalMAP (a continuation of PaCMAP) and DTNE.

**Reviewer 3**

This paper introduces PCC, a dimensionality reduction method aimed at preserving global structure while keeping competitive local structure. The idea is well motivated, and the biological imaging example is compelling, but the manuscript is not yet ready for publication. The main concerns are incomplete reporting and lack of statistical rigor. Page 9 refers to "Table ??" which is missing; the full benchmark results table with mean ± standard deviation must be included.

Experiments appear to be from single runs without error estimates; given the sensitivity of DR methods to initialization, results should be averaged across multiple seeds and supported by statistical tests.

> We thank the reviewer for this very valuable feedback. Following this feedback, we performed experiments over 10 random seeds, and extended the evaluation to focus on 20 biomedical datasets. This took significant compute time with several days to complete. We note that, interestingly, while the random seeds change the visual result, they have little impact on the evaluation metrics and therefore the seed std is negligible (below 1e-3 as shown in the figure). Therefore, when we report aggregated results over datasets we now report std over the 20 datasets, and when we show per dataset performance we showed the seed std.
>
> All our quantitative results now have error bars.

Methodological details are also thin: the reference point sampling strategy is unclear, the Spearman soft ranking approximation is not fully explained, and computational complexity is not discussed.

> Following this valuable feedback, we performed major revisions to the paper. For focus and simplification the Spearman objective was removed. We discuss computational complexity.

Hyperparameters such as K, β, and cluster numbers need justification or sensitivity analysis.

> We thank the reviewer for this feedback. Following this we added a dedicated ablation and hyper parameter sensitivity section showing the effect of the hyper parameters both visually and with quantitative metrics.

Minor issues include inconsistent notation, incomplete references, and figure captions that do not fully explain the results. The abstract could also be shortened to highlight the main contribution more clearly. Overall, I recommend major revisions. The method is interesting and shows promise, but the missing results and lack of statistical validation prevent proper evaluation at this stage. Once these issues are addressed, the work could be a strong contribution to the dimensionality reduction literature.

> We thank the reviewer for the very encouraging feedback. We performed a major revision, shortened the abstract, and tried to improve the figure captions.

**Reviewer 4**

This paper presents Preserving Clusters and Correlations (PCC), a novel dimensionality reduction (DR) method that achieves state-of-the-art global structure(GS) preservation while maintaining competitive local structure (LS) preservation. This paper somehow tried to demostrate its novelty with encouraging experimental results. However the following issues and suggestions are worth looking at comments

1. few phrases/sentences need restructuring (especially most sentences and paragraphs are very long) as well as some little grammatical errors such as listed below
   a. the section Introduction is spelt wrong "Introdution"
   b. in paragraph 2 of the introduction, there is a dot before and after [6], kindly correct
   c. in paragraph 3 of the introuction, "...close points in the clusters" can be revise to "...closely spaced points within clusters."
   d. under results/discussions, "...close the PCA" could be change to "...close to PCA"

   We thank the reviewer for this feedback, and fixed these as part of a major revision of the paper.

2. Some few problems in equations: the equation are not labeled, this is very difficult for references

   Following this feedback, all equations are now labeled.

3. previous works is too scanty thus several very closely related methods have been left out

   We thank the reviewer for this valuable feedback. We extended the previous work section. We also now elaborate more about the objectives of UMAP or PaCMAP, in order to discuss the relationship to them with attraction / repulsion terms in the correlation loss. We also mention related methods that used clustering as a signal for representation learning, as well as landmark methods in dimensionality reduction.

4. the figure labels contain too much text which is inappropriate

   We thank the reviewer for this feedback and significantly reduced the figure label text.

5. the section 6 titled discussion rather contains conclusion paragraphs

   We thank the reviewer for this valuable feedback. We have completely rewritten the discussion to focus on discussion.

6. under section 5. The full results are given in Table ??. several of these missing references exist in the document 7. in references a) Ref[2] ("???" year). b) Ref [16] publisher info incomplete ("???") as well as several inconsistent DOIs

We thank the reviewer for these and fixed these. We hope that now in general the formatting and clarity of the paper is greatly improved.
